# A rare human *TNFAIP3* variant reveals how A20 abundance is regulated by TAX1BP1

**DOI:** 10.64898/2026.03.25.714162

**Authors:** Cindy Eunhee Lee, Morgan Downes, Rochna Chand, Keisuke Horikawa, Vicki Athanasopoulos, Bahar Miraghazadeh, Matthew C. Cook

## Abstract

Variants in *TNFAIP3* (encoding A20) are among the most consistent non-MHC genetic associations with human autoimmune disease. Moreover, *TNFAIP3* haploinsufficiency confers an autosomal dominant autoinflammatory and autoimmune disease. Whilst these findings strongly suggest that quantitative changes in A20 predispose to pathology, there is currently no evidence that post-transcriptional modifications regulate A20 abundance. We used human reverse genetics approach to investigate this question. From a cohort of patients with complex immune diseases, we identified an ultrarare hypomorphic TNFAIP3 variant (A20^S254R^) that is both hypomorphic and prone to phosphorylation. One of five carriers of this variant was distinguished by autoimmunity and an NF-kB gain-of-function transcriptional signature. A search for modifiers identified a variant in *TAX1BP1* (p.Leu207Ile). We discovered that TAX1BP1 normally retrains A20 phosphorylation. Furthermore, TAX1BP1 preferentially binds phospho-A20, which explains the enhanced interaction between TAX1BP1^L207I^ and A20^S254R^. A20 phosphorylation regulates its abundance, not by MALT1-mediated degradation, but probably via autophagy. Thus, while A20 phosphorylation has been implicated in A20 protease activity, an epistatic interaction between rare human genetic variants reveals how phosphorylation also regulates A20 abundance.

## INTRODUCTION

Polymorphisms in *TNFAIP3* (encoding A20) are amongst the most consistent genome-wide associations with autoimmune disease, including systemic lupus erythematosus (SLE), rheumatoid arthritis, Sjogren’s syndrome, thyroiditis and multiple sclerosis (Ma & Malynn, 2012; Musone *et al*, 2008; Compagno *et al*, 2009; Bates *et al*, 2009; Ramos *et al*, 2011; Graham *et al*, 2008). Furthermore, some of these noncoding disease-associated *TNFAIP3* variants are expression quantitative trait loci, which suggests that NF-kB regulation is sensitive to small changes in A20 expression (Adrianto *et al*, 2011; Wang *et al*, 2013, 2016). Substantial reductions in A20 abundance are definitely pathological; *TNFAIP3* haploinsufficiency causes familial Behcet-like autoinflammatory syndrome-1 (AIFBL1; OMIM# 616774), which often presents with organ-specific autoimmunity, either alone or in combination with autoinflammatory manifestations (Aeschlimann *et al*, 2018a; Hori *et al*, 2019; Zhou *et al*, 2016; Takagi *et al*, 2017; Kadowaki *et al*, 2018).

In mice, complete A20 deficiency causes lethal, early-onset inflammation similar to that observed with human AIFBL1 (Lee *et al*, 2000). Numerous genetic manipulations in mice have probed the functional domains of A20 to explain this phenotype but have not yet yielded a conclusive account of the biochemical mechanism that links A20 to disease. The C-terminus of A20 contains seven zinc finger (ZF) domains that regulate E3 ligase activity, mono- and polyubiquitin binding, and protein interactions (Wertz *et al*, 2004; Boone *et al*, 2004; Komander & Barford, 2007; Bosanac *et al*, 2010). ZF7 has also been shown to mediate non-catalytic inhibition of IKKβ (Skaug *et al*, 2011). Mice with disruptions of ZF7 domains remain healthy to at least one year of age after which they develop psoriatic arthritis (Razani *et al*, 2020), while combined disruption of ZF4 and ZF7 results in accelerated inflammatory disease (De *et al*, 2014; Lu *et al*, 2013; Martens *et al*, 2020; Razani *et al*, 2020). A20 also contains an N-terminal ovarian tumour (OTU) domain in which deubiquitinating cysteine protease activity is mediated by a catalytic triad comprising Cys103, His256 and Asp70 that cleaves K11, K48 and K63 ubiquitin chains (Wertz *et al*, 2004; Boone *et al*, 2004). Remarkably, mice harbouring a specific deletion of the A20 OTU domain remain healthy (De *et al*, 2014; Lu *et al*, 2013). A20 is subject to phosphorylation at several sites and phosphorylation of Ser381 is thought to enhance A20 deubiquitinase activity (Wertz *et al*, 2015; Hutti *et al*, 2007). Putative phospho-defective alleles of *TNFAIP3* have been associated with autoimmune disease and attributed to reduced OTU activity (Zammit *et al*, 2019) but this mechanism alone is at odds with the benign consequences of complete OTU deletion.

Irrespective of the precise biochemical actions of A20 that lead to disease, reduction in A20 abundance is pathological. Furthermore, regulation of A20 abundance is particularly important in lymphocytes because they express high levels of A20 constitutively and after receptor ligation, A20 removal is probably necessary for efficient NF-kB activation (Lee *et al*, 2000; Catrysse *et al*, 2014; Coornaert *et al*, 2008; Tewari *et al*, 1995). Despite the importance of regulating A20 abundance, there remain large gaps in our understanding of how this is achieved. First, while there are two pathways for A20 disposal, proteolysis mediated by MALT1 paracaspase and p62-dependent autophagy (Kanayama *et al*, 2015; Coornaert *et al*, 2008) no post-transcriptional modifications have been identified that target A20 for disposal. Second, A20 has a number of binding partners, including ITCH, TAX1BP1, and TNIP1 (ABIN1) (Shembade *et al*, 2007; Heyninck *et al*, 1999; Shembade *et al*, 2008; Valck *et al*, 1999) that regulate assembly of A20-containing complexes but it remains unclear whether any of these interactions are affected by A20 phosphorylation, or whether they affect A20 disposal. Finally, there is little if any evidence of post-transcriptional regulation of A20 abundance in disease states.

In addition to loss-of-function variants in humans and knockout mouse models, rare damaging missense variants observed in humans offer a distinct opportunity to elucidate gene–gene interactions. Thus, we conducted a reverse genetics screen of human genomes for rare *TNFAIP3* variants and identified a candidate in two kindreds, although carriers were discordant for immune phenotypes. We identified a second variant in TAX1BP1, an interacting partner of A20. On further investigation we discovered that TAX1BP1 regulates A20 phosphorylation and that phosphorylation of A20 regulates its abundance.

## RESULTS

### Reverse genetics screen identifies a hypomorphic *TNFAIP3* variant underlying immune dysregulation

We screened a large cohort of human genomes from patients with undiagnosed immune diseases, including primary antibody deficiency and autoimmunity, for rare or novel variants in immune-related genes. We identified a *TNFAIP3* variant (c.762C>A; p.Ser254Arg) that is ultrarare (MAF<0.00002, GnomADv4.1.0) in five individuals from two unrelated kindreds (**Fig. 1a**). Ser254 is highly conserved across species expressing A20 (**Fig. 1b**), is located in the OTU domain, and is predicted to be damaging by *in silico* tools (CADD=22.8; Polyphen2=0.998; SIFT=0.09). p.Ser254Arg is classified as likely pathogenic by AlphaMissense (Cheng *et al*, 2023) (**Fig. 1c**). Ser254 is located on the β4 sheet of the OTU catalytic core, in close proximity to the proteolytic catalytic triad (Cys103, His256, Asp70) (**Fig. 1d-e; Supplementary Fig. 1**) (Komander & Barford, 2007).

**Figure 1.**
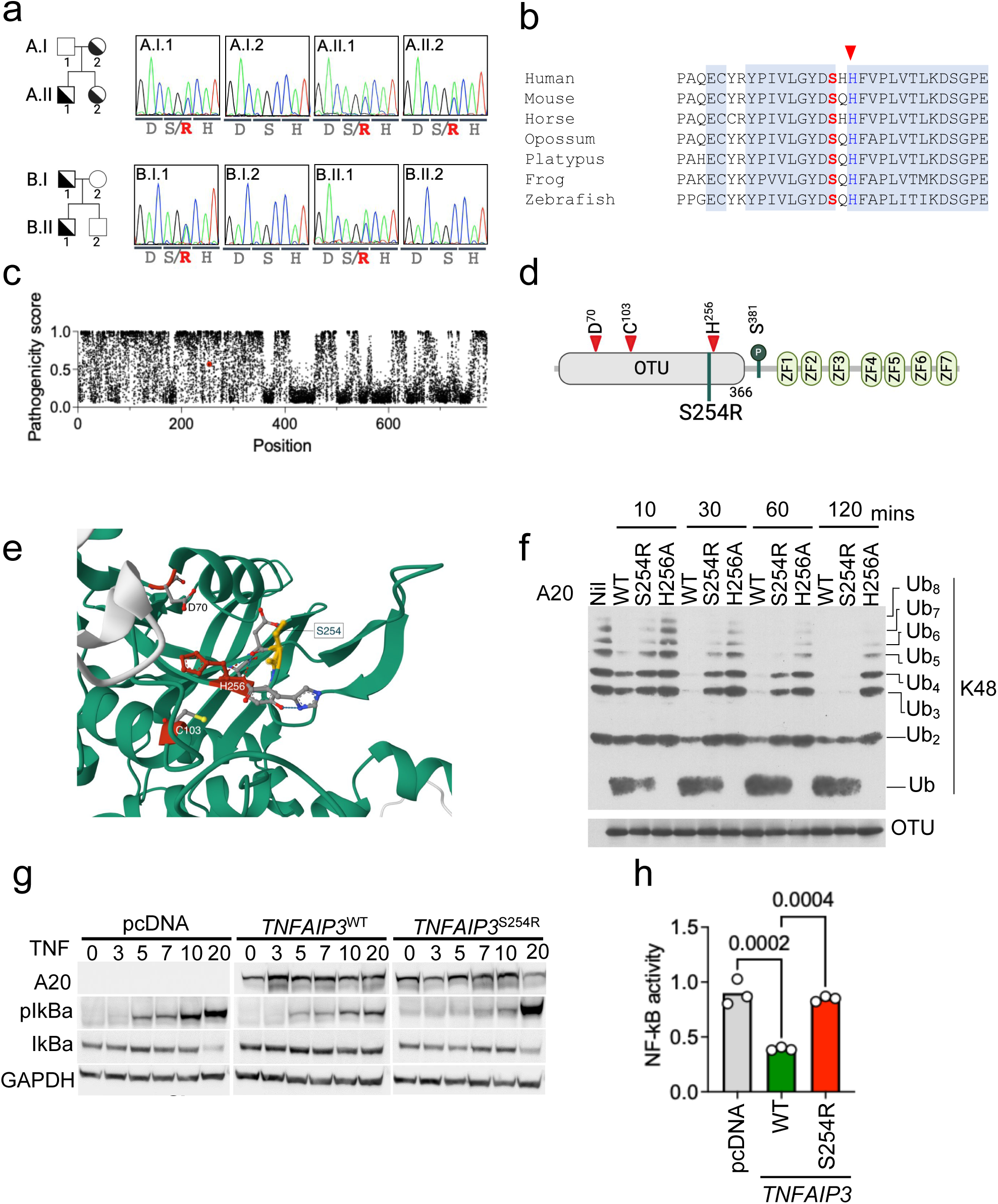
Function of A20^S254R^. **a**. Pedigrees of two unrelated kindreds indicating carriers of *TNFAIP3*^S254R^ (semi-filled symbols), confirmed by Sanger sequencing. **b**. Evolutionary conservation of Ser254. His256 (red arrow) is part of the OTU catalytic triad. **c**. AlphaMissense pathogenicity scores for *TNAFAIP3*. S254R, red symbol. **d.** Schematic of human A20 indicating ovarian tumour (OTU) domain and seven zinc finger (ZF) domains. Residues comprising the OTU catalytic triad (red arrows), phosphorylation site (Ser381) and missense mutation (Ser254) are highlighted. **e**. Ribbon diagram of A20 structure to show proximity of Ser254 to Cys103 and His256 (AlphaFold2). **f**. Relative abundance of various K48-linked polyubiquitin chains after incubation with purified A20^S254R^, A20^WT^ or A20^H256A^ OTU constructs (loaded as shown). At indicated times, the enzyme reaction was stopped and hydrolysis of polyubiquitin chains was examined by Western blotting. **g**. Western blot analysis of IkBα and phosphorylated IkBα (pIkBa) after TNF stimulation of HEK293T transfected with *TNFAIP3*^WT^ or *TNFAIP3*^S254R^ constructs. **h**. NF-kB activity determined by luciferase. Constructs for empty vector, wildtype A20 or S254R A20 were co-transfected with Nifty and Renilla into HEK293. NFkB activity (Nifty) was normalised with Renilla activity.

Substitution of Ser254 with arginine (A20^S254R^) is predicted to result in steric hindrance of OTU substrates and impact deubiquitination activity. To test this prediction, we expressed and purified wild-type (WT) and A20^S254R^-derived OTU domains, plus a known catalytically inactive OTU domain (OTU^H256A^) and incubated them with K48-linked polyubiquitin chains. OTU^WT^ cleaved most of the polyubiquitin chains into monoubiquitin (Ub), whereas OTU^S254R^ exhibited reduced cleavage, most evident at early timepoints, with an effect intermediate between OTU^WT^ and catalytically inactive OTU^H256A^ (**Fig. 1f**). These findings demonstrate that the OTU^S254R^ domain has impaired catalytic activity compared to the wild-type OTU domain. To determine whether A20^S254R^ modulates NF-kB activity, we transiently transfected HEK293 cells with *TNFAIP3*^WT^ or *TNFAIP3*^S254R^ constructs. After stimulation with TNF, *TNFAIP3*^S254R^ transfectants exhibited enhanced NF-kB activation, indicated by increased pIkBα and reduced degradation of IkBα compared to *TNFAIP3*^WT^ transfectants after stimulation (**Fig. 1g**). We used the same system co-transfected with a luciferase readout of binding to the NF-kB response element and confirmed that A20^S254R^ exhibits reduced NF-kB inhibitory function compared to A20^WT^ (**Fig. 1h**). Taken together, these findings demonstrate that A20^S254R^ is a catalytically hypomorphic variant that disrupts A20-mediated control of NF-κB signalling.

### Heterogenous clinical and cellular phenotypes associated with A20^S254R^

In kindred A, one of the carriers of the *TNFAIP3*^S254R^ variant (A.II.1) presented with panhypogammaglobulinaemia and autoimmune cytopenias that required splenectomy, and was diagnosed with common variable immune deficiency (CVID). The same *TNFAIP3* variant was also identified in his sibling (A.II.2), who is IgA deficient, and his mother (A.I.2), who has multiple sclerosis and normal antibody levels. In kindred B, one of the carriers (B.II.1) presented with recurrent respiratory tract infections and was diagnosed with CVID but had no history of autoimmunity. The other carrier (B.I.1) has rheumatoid arthritis and normal serum immunoglobulin levels.

We analysed the peripheral lymphocytes from both kindreds and included an unrelated splenectomised patient as an additional control. Notably, we identified B- and T-cell phenotypes in A.II.1 that were not found in any other A20^S254R^ carriers, or in the splenectomised control (**Fig. 2a, Supplementary Fig. 2a-c**). B cell analysis revealed a reduction in naïve and conventional memory cells, and relative expansions of transitional B cells and CD21^low^ atypical memory B cells in A.II.1. Similarly, the T cell compartment revealed relatively few naïve T cells and a high proportion of memory T cells (**Fig. 2a**), with a distinct effector T cell distribution and high proportion of exhaustion markers in A.II.1 (**Supplementary Fig. 2d**).

**Figure 2.**
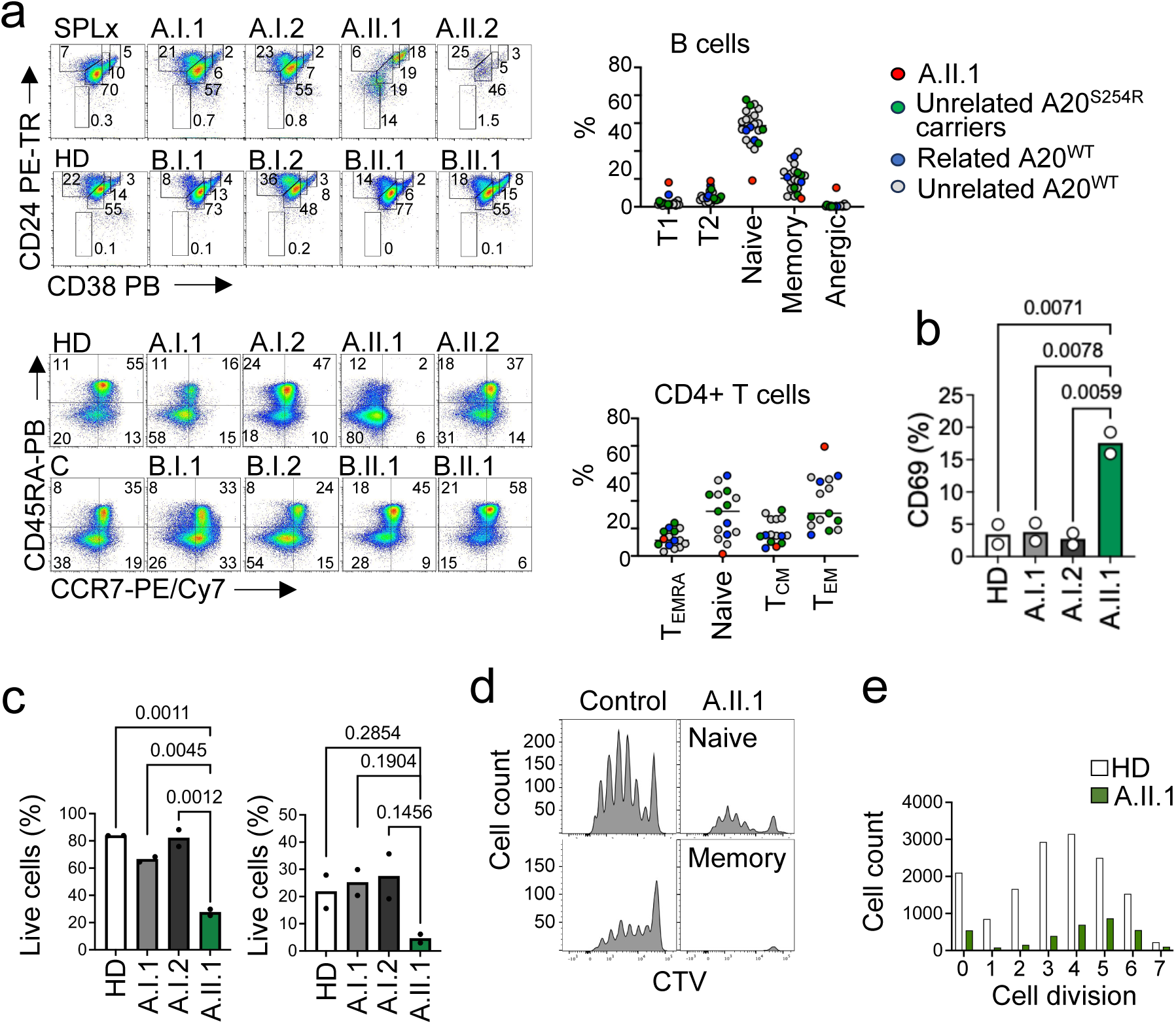
Lymphocyte function and phenotype. **a**. Representative flow cytometric analysis of PBMCs from indicated kindred members and an unrelated splenectomised control (SPLx). B cells were gated on CD19^+^ CD3^-^. Region percentages are shown for naïve B cells (CD24^+^, CD38^+^), transitional stage 1 B cells (CD24^+++^, CD38^+++^), transitional stage 2 B cells (CD24^++^, CD38^++^), memory B cells (CD24^++^, CD38^+^), atypical memory B cells (CD24^-^, CD38^-^), and plasma cells (CD24^-^CD38^+++^). CD4+ T cells were gated on CD3+ CD4+ CD19-. Region percentages are shown for CD45RA+ effector memory (T_EMRA_, CD45RA^+^, CCR7^-^), naïve (CD45RA^+^, CCR7^+^), central memory (T_CM_, CD45RA^-^, CCR7^-^), and effector memory T cells (T_EM_, CD45RA^-^, CCR7^+^). Summary plot of B and T cell subsets (right panels). **b.** Summary plot of CD69 expression on unstimulated (ex vivo) naïve T cells from indicated donors, from 2 independent experiments. **c.** Summary of viability of naïve T cells with (right panel) or without (left panel) T cell activation, from 2 independent experiments. **d**. Proliferation assay. Sorted naïve T cells were labelled with cell trace violet and stimulated with anti CD2,CD3, CD28 for 5 days and the total cell number and CTV diluted cells were assessed. **e.** Bar graph of viable cell counts by division number.

Next, we investigated plasmablast formation. B cells were stimulated with IL-21 and CpG for five days. B cells from A.II.1 did not form plasmablasts while normal plasmablast differentiation was observed in B cells from healthy controls and other A20^S254R^ carriers (**Supplementary Fig. 3**). Functional analysis of circulating T lymphocytes was performed on sorted populations of naive CD4+ cells and effector memory T cells, which we stimulated in vitro with CD2/CD3/CD28 coated beads for 24 hours. Approximately 15% of naïve T cells from A.II.1 were CD69 positive even in the absence of in vitro stimulation, which represents a significant difference from controls and other *TNFAIP3*^S254R^ carriers (**Fig. 2b, Supplementary Fig. 4**). During in vitro lymphocyte analysis, we noted a disproportionate loss of viability in cells from A.II.1. In the absence of stimulation, approximately 70% of naïve T cells from A.II.1 were non-viable compared with 10-20% in controls and other A20^S254R^ carriers. After stimulation, the cell loss from A.II.1 was even more pronounced (**Fig. 2c**). To explore this further, we enumerated live T cells during proliferation after stimulation. Sorted naïve T cells were labelled with Cell Trace Violet (CTV) and stimulated with anti-CD2/CD3/CD28 for 5 days. While proliferation of T cells from all donors was similar, total cell numbers from cultures of A.II.1 cells were significantly reduced compared with those from controls (**Fig. 2d-e**). A similar defect in cell survival despite normal proliferation defects was previously reported in mice with a T cell-specific deletion of *Tnfaip3* (Onizawa *et al*, 2015). Taken together, compared with other carriers of hypomorphic A20^S254R^, A.II.1 exhibits a distinct clinical phenotype characterised by severe haematological autoimmunity and generalised lymphadenopathy, and more profound alterations in immune cell phenotype and function.

### *TAX1BP1^L307I^* is a potential modifier of *TNFAIP3*

So far, our data demonstrate that A20^S254R^ is catalytically hypomorphic and correlates with increased NF-κB activity, as well as clinical and cellular abnormalities that are plausibly attributable to dysregulated NF-κB signalling. The exceptional biochemical and cellular phenotype observed suggests the existence of a genetic modifier of *TNFAIP3*^S254R^ in the proband. We investigated this hypothesis with the aim of identifying a novel epistatic interaction with *TNFAIP3* that might contribute to A20 regulation.

To investigate whether the markedly dysregulated immune phenotype in A.II.1 was driven by altered NF-κB signalling, we examined lymphocyte transcriptomes from A.II.1. To avoid the risk of differences in lymphocyte differentiation confounding our functional analysis, we sorted naïve B cells (CD19+ CD21+ CD10- CD27-) and T cells (CD3+ CD4+ CD45RO-) from A.II.1 and healthy controls then analysed their transcriptomes after stimulation with anti-IgM and anti-CD2/3/28, respectively. This revealed substantial upregulation of NF-kB-responsive genes in A.II.1 compared with controls (**Fig. 3a-b; Supplementary Fig. 5**). *IL8* and *CCL2* (MCP-1) were highly expressed in both naïve T and B cells from A.II.1. In addition, *TNFAIP3* itself, which is an NF-kB target gene, was also noted to be expressed at high levels in both B and T cells from A.II.1 (**Fig. 3a**). These results are reminiscent of previous findings from both *Tnfaip3*^C103A^ and *Tnfaip3*^ZF4^ mice, where *Il1, Tnf, Ccl2* and *Tnfaip3* were all reported to be upregulated (Lu *et al*, 2013).

**Figure 3.**
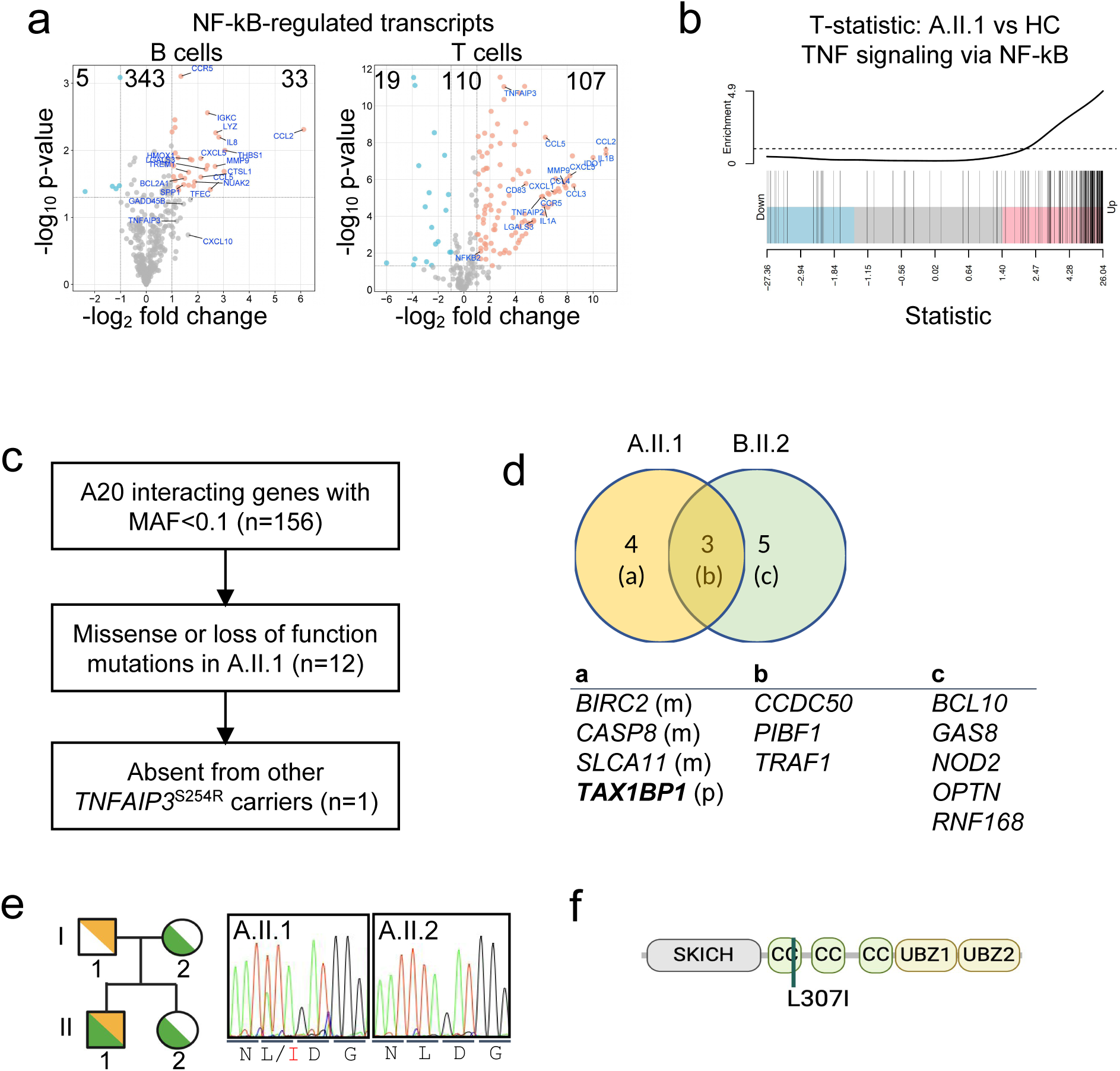
*TAX1BP1*^L307I^ is a putative genetic modifier of *TNFAIP3*^S254R^. **a**. Volcano plots NF-kB target gene expression determined by bulk RNA sequencing of sorted naïve B cells (left panel) or sorted naïve T cells (right panel) from A20^S254R^ carrier (A.II.1) or unrelated A20^WT^ controls (C, C1, C2) stimulated with anti-Ig or anti CD2,CD3 andCD28, respectively. **b**. GSEA analysis of NF-kB target gene expression. Data derived from three technical replicates. **c**. Summary of filtering strategy used to identify potential modifiers of *TNFAIP3*^S254R^ in A.II.1. **d**. Candidate genes (containing variants) from kindreds A and B are shown. Genes with variants unique to A.II.1 are shown in column (a) (right panel) with inheritance shown (m, maternal; p, paternal). **e**. Kindred A indicating transmission of *TAX1BP1*^L307I^ (orange) and *TNFAIP3*^S254R^ (green) variants, validated by Sanger sequencing (right panel) **f.** Schematic representation of human TAX1BP1 domain structure. SKICH, SKIP carboxyl homology; CC, coiled-coil; UBZ, ubiquitin binding zone. Leu^307^ is indicated.

We next explored potential epistatic interactions affecting A20 regulation. We began by interrogating variants in the A.II.1 genome affecting genes known to encode proteins that interact with A20. String (Szklarczyk *et al*, 2015) and BioGrid (Oughtred *et al*, 2021) searches identified 156 genes that interact with *TNFAIP3*; 12 of these genes harboured nonsynonymous or splice-site mutations in the A.II.1 genome (**Fig. 3c**). We then excluded those variants *in cis* with maternally transmitted *TNFAIP3*^S254R^ in A.II.1. This left one candidate, a heterozygous missense variant in *TAX1BP1* (c.919T>A; p.Leu307Ile) that was transmitted paternally to A.II.1 (**Fig. 3d-e**). The TAX1BP1^L307I^ variant is located in the coiled-coil domain, which contributes to dimerization, binding to TRAF6, and protein-protein interactions important for autophagy (Ling & Goeddel, 2000; Zhang *et al*, 2024; Hu *et al*, 2018) (**Fig. 3f, Supplementary Fig. 6**). Although *TAX1BP1* c.919T>A is not rare (MAF = 0.10, GnomADv4.1.0), Leu307 is highly conserved and *in silico* tests predict that substitution with isoleucine is possibly damaging (Polyphen2=0.278; SIFT=0.78, CADD=19.9).

TAX1BP1 functions as a ubiquitin-binding adaptor that brings A20 into proximity with substrates such as Ubc13, UbcH5c, and the E3 ligase TRAF6, thereby facilitating negative regulation of NF-κB signalling (Shembade et al., 2010; Ling & Goeddel, 2000; Valck et al., 1999). In addition to this adaptor role, TAX1BP1 also acts as a selective autophagy receptor that recognises ubiquitinated cargo and targets it for degradation via the autophagy-lysosome pathway (White et al., 2023). Here, we identified TAX1BP1 as a compelling candidate genetic modifier of TNFAIP3, providing a plausible mechanistic basis for epistatic regulation of A20 functions and NF-κB signalling in A.II.1.

### A20^S254R^ results in hyperphosphorylation, leading disposal of A20

To investigate putative epistasis between the *TNFAIP3* and *TAX1BP1* variants in lymphocytes, we generated a *TNFAIP3* and *TAX1BP1* double-deficient Raji B cell line (DKO) by CRISPR/Cas9 gene editing (**Fig. 4a**; **Supplementary Fig. 7**). DKO cells expressed higher levels of CD69 compared to wildtype (**Fig. 4b**). These findings were consistent with the enhanced CD69 expression observed on naïve T cells from A.II.1 (**Fig. 2b**).

**Figure 4.**
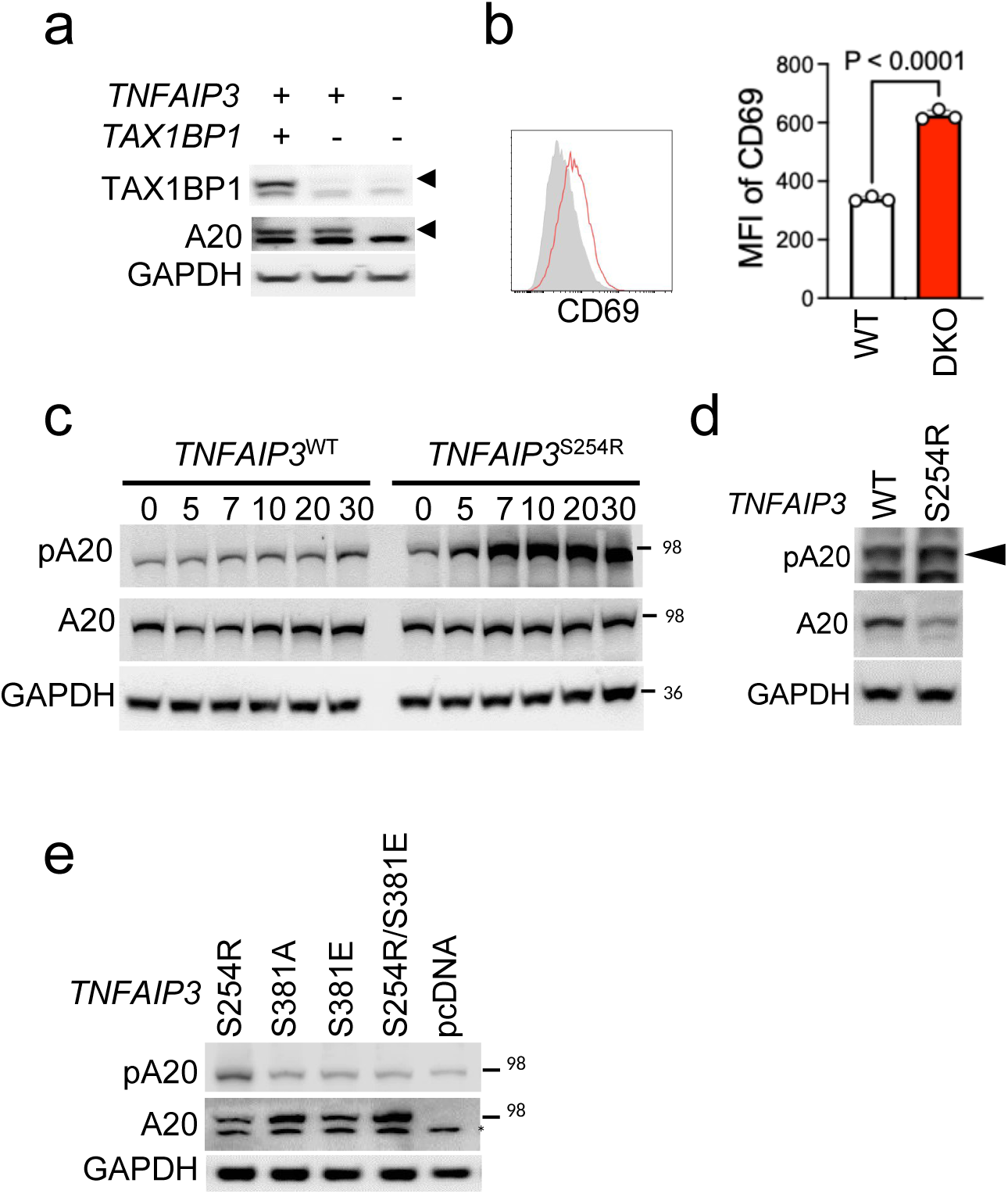
Phosphorylation and cleavage of A20. **a**. Western blot analysis of indicated cell lines. **b**. Flow cytometry histogram of CD69 expression by WT (grey), TAX1BP1-deficient (green) or *TAX1BP1/TNFAIP3*-double deficient (DKO, red) Raji cells with summary of mean fluorescence intensities (MFI) of CD69 expression by indicated Raji cells from three independent experiments. **c**. Western blot comparing phosphorylation of A20^WT^ and A20^S254R^ in HEK293T transfectants after stimulation with TNF. **d**. Comparison of pA20 in the presence or absence of TAX1BP1. **e.** Abundance of A20 and pA20 was assessed after indicated phospho-defective or phospho-prone *TNFAIP3* variants were transfected into DKO Raji cells.

To explore the functional consequences of the *TNFAIP3*^254R^ variant, we first assessed A20 activity. Since A20 phosphorylation has been reported to regulate its catalytic activity, and autoimmune-prone variants frequently exhibit defects in phosphorylation (Zammit et al., 2019; Wertz et al., 2004; Hutti et al., 2007), we examined whether A20^S254R^ displays impaired phosphorylation that could account for defective NF-κB inhibition. HEK293 cells were transiently transfected with *TNFAIP3*^WT^ or *TNFAIP3*^S254R^ constructs and stimulated with a range of TNF concentrations to assess phospho-A20 (pA20). Surprisingly, A20^S254R^ exhibited elevated Ser381 phosphorylation at baseline and a marked increase following TNF stimulation compared with WT A20 (**Fig. 4c**). Unlike most cell types, lymphocytes constitutively express A20, suggesting that A20 regulation may differ in this cellular context. To determine whether A20^S254R^ also exhibits increased phosphorylation in lymphoid cells, *TNFAIP3* and *TAX1BP1* double-deficient Raji B cell line (DKO) were transiently transfected with *TNFAIP3*^WT^ or *TNFAIP3*^S254R^ constructs and pA20 levels were assessed. After accounting for the lower expression of total A20, A20^S254R^ exhibited elevated Ser381 phosphorylation (especially as a proportion of total A20) compared to A20^WT^ (**Fig. 4d**).

As outlined above, tight regulation of A20 abundance is important for maintaining normal immunity. Since total A20^S254R^ protein levels were modestly reduced compared with A20^WT^, we investigated the potential link between A20 phosphorylation and protein abundance. In order to test whether A20 turnover is phospho-dependent, we took advantage of the phospho-prone *TNFAIP3*^S254R^ mutant, which we transiently expressed in *TNFAIP3*/*TAX1BP1* DKO Raji B cells. We compared A20^S254R^ abundance with a phosphorylation-defective protein (A20^S381A^) and a version of the protein (A20^S381E^) that functionally mimics constitutive phosphorylation (although it is not recognised by the anti-pA20 antibody). Abundance of total A20^S254R^ was similar to A20^S381E^, while there was a relative increase in abundance of phospho-defective A20^S381A^ (**Fig. 4e**). We also engineered a double mutant that is phospho-defective at Ser381 (*TNFAIP3*^S254R/S381A^). This mutant resulted in a further increase in A20 protein compared with A20^S254R^, indicating that turnover of A20^S254R^ is phosphorylation-dependent (**Fig. 4e**).

### TAX1BP1 regulates pA20

Our results indicate that A20^S254R^ undergoes hyperphosphorylation that results in reduced A20 abundance. We investigated whether the phenomenon occurs in patient cells. We stimulated PBMCs from A.II.1 and two unrelated healthy donors with PMA/ionomycin, then immunoblotted for total A20 and pA20. A20 abundance in cells from A.II.1 was reduced in unstimulated PBMCs relative to healthy donors (HD) and declined further after TNF stimulation (**Fig. 5a-b**). The decrease was observed as early as two minutes of stimulation, whereas subtle changes in A20 abundance were observed in control samples, suggesting the potential epistatic effects of the two missense variants A20^S254R^ and TAX1BP1^L307I^ identified in patient A.II.1. *TNFAIP3* is transcriptionally regulated by NF-kB, and consistent with this, we observed high *TNFAIP3* expression by RNAseq and microarray (**Fig. 3a**). Given the abundance of *TNFAIP3* mRNA, the observed reduction in A20 protein levels is therefore likely to be post-transcriptional.

**Figure 5.**
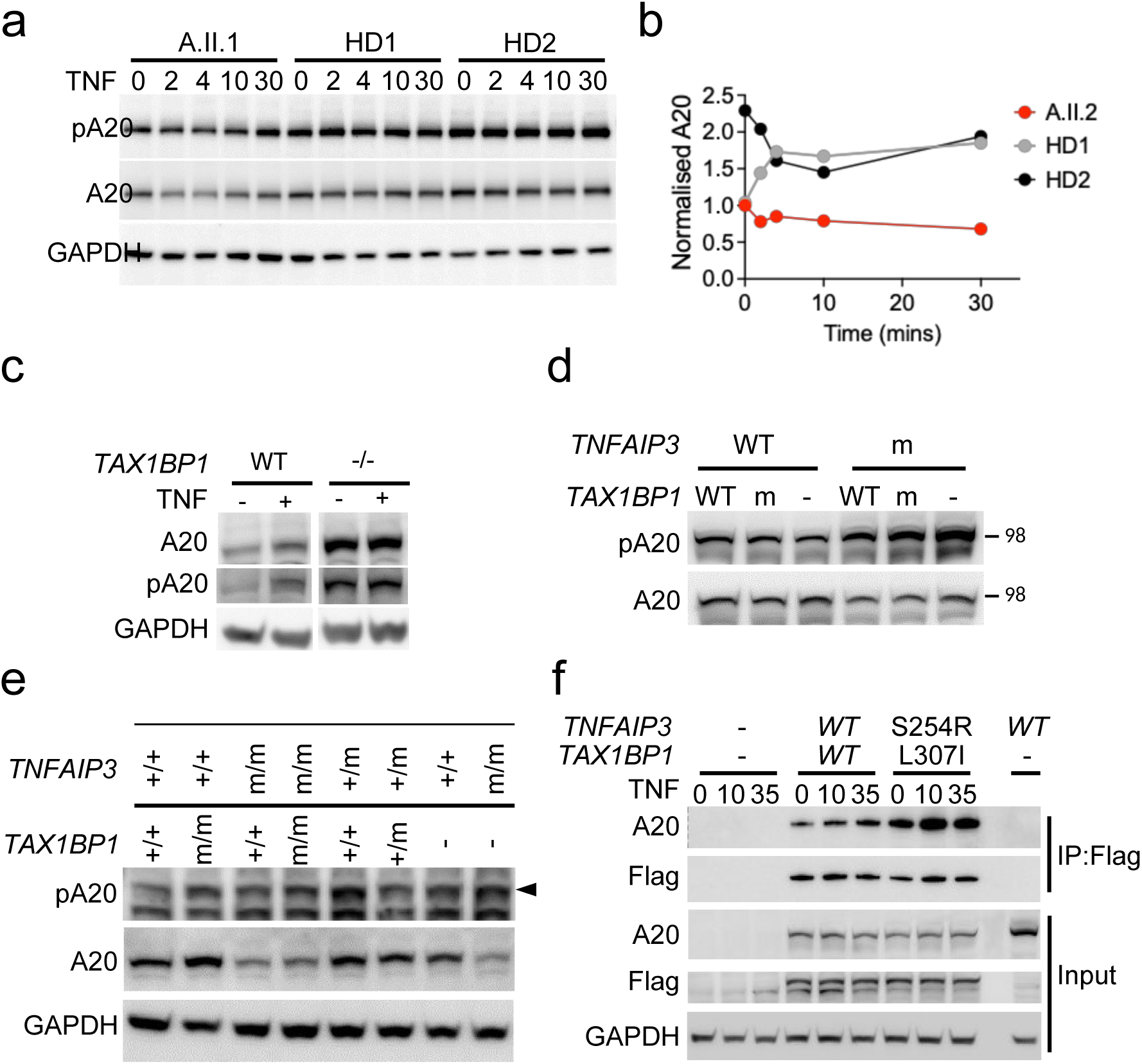
TAX1BP1-dependent degradation of pA20. **a-b.** Western blot of A20, pA20 and pIKKαβ from human PBMCs in the presence or absence of stimulation with TNF for the indicated times, with quantification (**b**). **c**. Analysis of abundance of A20 and pA20 and NF-kB activation in wild type or *TAX1BP1^-^*^/-^ Raji cells in the presence or absence of TNF stimulation. **d**-**e.** Abundance of A20 and pA20 after co-transfection of indicated combinations of *TAX1BP1*^WT^ or *TAX1BP1*^L307I^ and *TNFAIP3*^WT^ or *TNFAIP3*^S254R^ into HEK293 cells (**d**) or Raji cells (**e**). **f**. Co-immunoprecipitation of A20^WT^ and Flag-TAX1BP1^WT^ or A20^S254R^ and Flag-TAX1BP1^L307I^ after transfection into HEK293. TAX1BP1 was pulled down with anti-Flag.

We postulated that TAX1BP1 is involved in A20 phosphorylation. To investigate this, we generated a *TAX1BP1-*deficient Raji B cell line by CRISPR/Cas9 gene editing and examined the role of TAX1BP1 in modulating A20 phosphorylation. Analysis of *TAX1BP1*^-/-^ cells showed an increase in pA20 in resting status to levels similar to those seen after activation of TAX1BP1 WT cells (**Fig. 5c**). pA20 was increased at baseline and after TNF stimulation in the absence of TAX1BP1.

Next, we investigated the potential epistatic effects of the two missense variants A20^S254R^ and TAX1BP1^L307I^ identified in patient A.II.1. We co-expressed *TNFAIP3*^WT^ or *TNFAIP3*^S254R^ with *TAX1BP1*^WT^ or *TAX1BP1*^L307I^ in HEK293 cells. We observed that TAX1BP1^WT^ reduced the abundance of pA20 observed with the A20^S254R^ variant when compared with TAX1BP1^L307I^ or absence of TAX1BP1 (**Fig. 5d**). To explore TAX1BP1 functions in lymphocytes, we performed similar experiments in transfected Raji B cells. Again, we observed increased pA20 in the absence of TAX1BP1 or in the presence of TAX1BP1^L307I^ (**Fig. 5e**). This result suggests that increased pA20 conferred by A20^S254R^ increases NF-kB activity via A20 insufficiency, and pA20 was regulated by TAX1BP1.

To account for these findings, we postulated that pA20 (and therefore A20^S254R^) might be unusually partitioned towards the TAX1BP1 pathway for A20 disposal; therefore, we tested the physical interactions of mutant and WT versions of A20 with TAX1BP1. Immunoprecipitation of Flag-tagged TAX1BP1 pulled down more A20^S254R^ than A20^WT^, consistent with an increased propensity for A20^S254R^ to interact with TAX1BP1^L307I^ (**Fig. 5f**).

### A20 disposal

To test whether the A20 turnover is mediated by cleavage via the MALT1 paracaspase pathway (Coornaert et al., 2008), we transfected HEK293T cells with *TNFAIP3, MALT1,* and *BCL10* in the presence or absence of *TAX1BP1*. A20 was indeed cleaved by MALT1 with BCL10, however, additional TAX1BP1 did not affect the efficiency of this cleavage (**Fig. 6a**). To further explore this mechanism, we investigated interactions among MALT, TAX1BP1, and A20 by immunoprecipitation. The results demonstrated a strong interaction between TAX1BP1 and A20, but no interaction between TAX1BP1 and MALT1(**Fig. 6b**), which argues against participation of TAX1BP1 in MALT1-mediated A20 cleavage. Since MALT1 paracaspase activity is normally inhibited by TRAF6 (O’Neill *et al*, 2021), and TAX1BP1 serves as an adaptor for TRAF6 interaction with A20 (Shembade *et al*, 2007), we then tested the possibility that TAX1BP1 partitions A20 for degradation by the MALT1-TRAF6 complex. HEK293T cells were co-transfected with *TNFAIP3*, *MALT1*, and *BCL10* with escalating doses of *TRAF6*. As anticipated, the addition of *TRAF6* resulted in a dose-dependent reduction in A20 cleavage by MALT1 (**Fig. 6c**, lanes 1-3). In contrast, when cells were transfected with a fixed dose of TRAF6 and increasing amounts of TAX1BP1, no change in MALT1-mediated A20 cleavage was observed (**Fig. 6c**, lanes 4-9). However, a dose-dependent reduction in full-length A20 was detected (**Fig. 6c**, lanes 4-9), suggesting that TAX1BP1 promotes A20 degradation independently of MALT1-mediated cleavage.

**Fig. 6.**
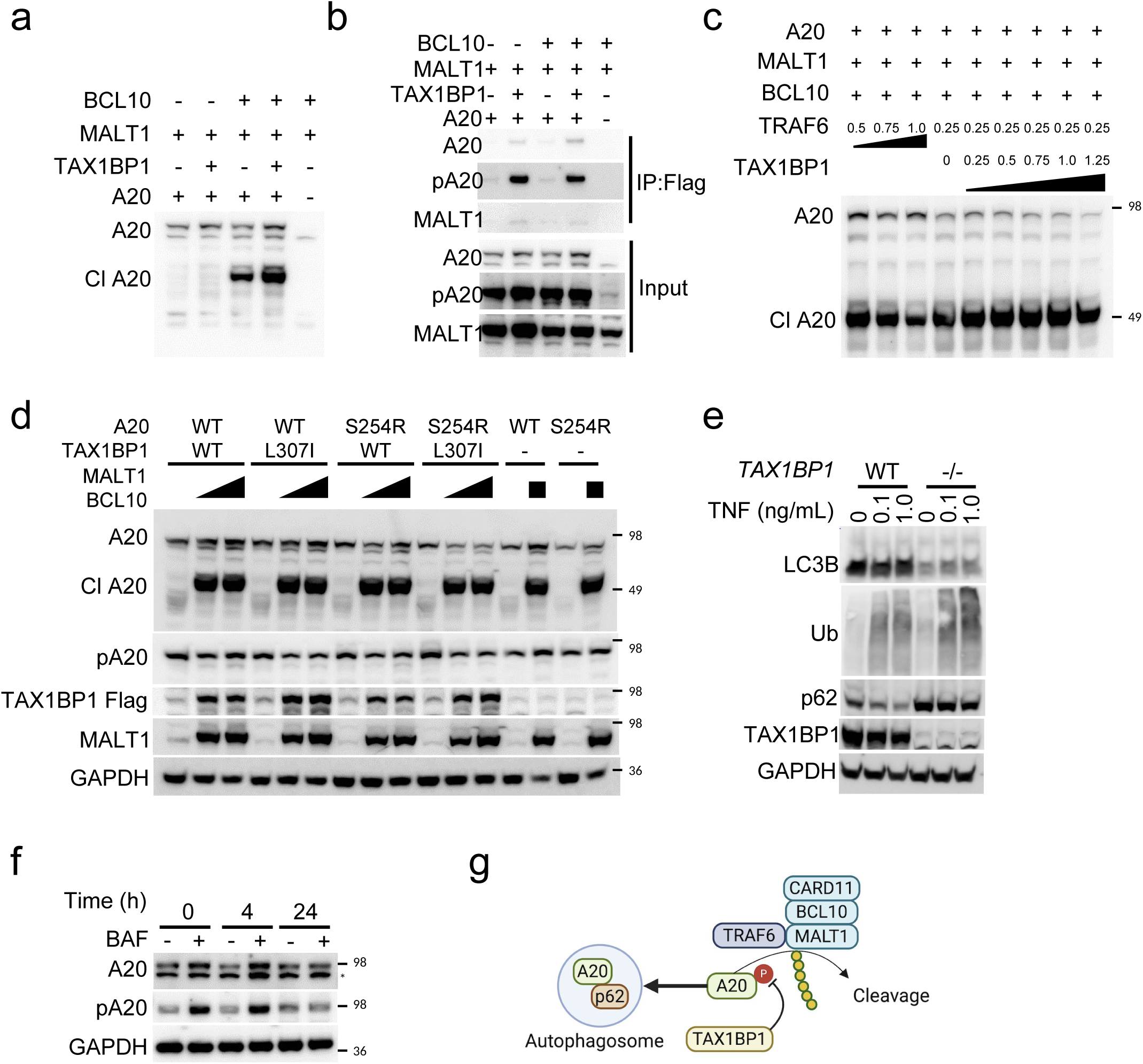
Selective TAX1BP1-mediated autophagy of phosphorylated A20. **a.** Western blot analysis of A20 cleavage in the presence of BCL10, MALT1 and TAX1BP1. **b.** Interaction between A20 and MALT1. TAX1BP1 was pulled down with anti-Flag and MALT1 and A20 were assessed by Western blot. **c**. Western blot of A20 cleavage in the presence of graded amounts of TRAF6 and TAX1BP1. *TNFAIP3, MALT1, BCL10* and *TRAF6* were transfected with or without *TAX1BP1* into HEK293. **d**. Analysis of MALT1-mediated A20 cleavage and TAX1BP1-mediated A20 degradation. *TNFAIP3^WT^* or *TNFAIP3*^S254R^ were co transfected with *TAX1BP1*^WT^ or *TAX1BP1*^L307I^ into HEK293 in the presence or absence of different amounts of MALT1 and BCL10. **e**. TAX1BP1 regulates autophagy. *TAX1BP1*^WT^ or *TAX1BP1*^-/-^ cells were stimulated with TNF. Whole cell lysates immunoblotted for LC3B, Ubiquitin (Ub) and p62. **f.** Abundance A20 and pA20 were assessed after culturing in glutamine free media +/- bafilomycin (BAF). **g**. Model summarising interaction between TAX1BP1 and pA20 (see text).

To compare A20 S254R and wild-type A20 in the presence of wild-type TAX1BP1 or the TAX1BP1^L307I^ variant, we assessed their co-expression under the same conditions. A20 cleavage was unchanged, however, full-length A20 levels differed between conditions. There was no reduction of wild-type A20 in the presence of TAX1BP1, whereas the A20^S254R^ variant exhibited a marked decrease in the presence of wild-type TAX1BP1. This reduction was more pronounced in the presence of TAX1BP1^L307I^, indicating increased susceptibility of the A20^S254R^ variant to TAX1BP1-associated degradation (**Fig. 6d**). Phospho-defective versions of A20 readily undergo cleavage (**Supplementary Fig. 8**).

A20 has also been reported to undergo autophagosomal degradation in a p62-dependent manner (Kanayama et al., 2015). To evaluate this mechanism, we analysed TAX1BP1-deficient cells. As expected, TAX1BP1 deficiency was associated with increased p62 levels and reduced LC3 abundance (**Fig. 6e**). TAX1BP1 functions as a cargo receptor that associates with NBR1-positive p62–ubiquitin condensates and promotes recruitment of the scaffold protein FIP200, a key step in autophagosome formation and selective autophagy (Turco et al., 2021). To induce autophagy in a TAX1BP1-independent manner, we subjected cells to nutrient starvation and examined pA20 and total A20 levels. Nutrient starvation increased A20 turnover, and this effect was abrogated by treatment with the autophagy inhibitor, bafilomycin A1 (F**ig. 6f**). These findings suggest that phosphorylation-dependent A20 turnover is mediated, at least in part, by autophagy.

## DISCUSSION

We have identified a mechanism of post-transcriptional regulation of A20 abundance. TAX1BP1 controls A20 phosphorylation at Ser381, and phosphorylation of A20 contributes to its disposal. TAX1BP1 is a known binding partner of A20 and is an established facilitator of autophagy for NF-kB regulation but its specific function in A20 phosphorylation and regulation of A20 abundance has not been appreciated until now (Shembade *et al*, 2007, 2010; White *et al*, 2023; Valck *et al*, 1999). Furthermore, phosphorylation of A20 has been implicated as an enhancer of its catalytic activity. Our findings now indicate that phosphorylation also regulates A20 abundance (Wertz *et al*, 2015; Hutti *et al*, 2007). This may explain why relatively few studies have directly tested A20 phosphorylation in vivo, despite numerous in vitro studies demonstrating the functional significance of phosphorylated A20.

We employed a reverse genetic approach to identify a novel *TNFAIP3* variant, which emerged as particularly informative because A20^S245R^ is intrinsically phospho-prone and catalytically hypomorphic. The reduction in A20 abundance conferred by its enhanced phosphorylation explains the paradoxical increase in NF-kB activity despite enhanced phosphorylation. Characterisation of a variant in the A20 interacting partner TAX1BP1 was also fortuitous, because this led us to discover that absence of TAX1BP1 results in enhanced phosphorylation of WT A20 (**Fig. 6g**). Thus, TAX1BP1 helps to maintain total A20 for IKK inhibition.

Previously, A20^S381A^ was shown to both attenuate cleavage of K63-linked tetraubiquitin by the OTU domain and enhance ubiquitination by the ZnF4 domain through an unknown mechanism (Wertz *et al*, 2015). It is, however, difficult to explain the association between pA20 and disease by this mechanism alone, as mice lacking either OTU or ZF4 do not recapitulate the A20 deficiency phenotype. Our findings and those previously reported mechanisms are not mutually exclusive, and a reduction in A20 abundance as a result of its phosphorylation provides a plausible mechanism of disease.

There are two pathways for the disposal of total A20. MALT1 paracaspase-mediated cleavage, which we confirm is dependent on BCL10 and is negatively regulated by TRAF6 (O’Neill *et al*, 2021), and autophagy. Our results demonstrate that phosphorylation-dependent A20 turnover does not result from MALT1-mediated cleavage. On the other hand, we show that specific disposal of pA20 is autophagy-dependent. Our results reveal that in the presence of MALT1 and BCL10, WT TAX1BP1 results in a dose-dependent reduction in A20 abundance. Furthermore, disposal of pA20 is amplified by the TAX1BP1^L307I^ variant, which exhibits enhanced binding of A20^S254R^. Previously, TAX1BP1 was shown to promote autophagy to regulate the availability of L-cysteine during T cell activation (Whang *et al*, 2017). Taken together, our findings are consistent with a model in which TAX1BP1 binds preferentially to pA20 to partition it for autophagy.

TAX1BP1 emerges as an important regulator of T cell function therefore abundance of TAX1BP1 is likely to contribute to fine tuning of A20 activity. Indeed, we show that MALT1 and BCL10 results in an increase in TAX1BP1. Regulation of A20 by TAX1BP1 is likely to be important in cells that express BCL10. In particular, lymphocytes express high levels of A20 and BCL10, and A20 disposal has been postulated to be a prerequisite for NF-kB activation (Catrysse *et al*, 2014; Coornaert *et al*, 2008). This complex dynamic interplay might explain the diversity in reported NF-kB phenotypes conferred by TAX1BP1 deficiency and mutation (Shembade *et al*, 2007; Iha *et al*, 2008; Whang *et al*, 2017; Morriswood *et al*, 2007). Further work will be required to determine how different stimuli in various cell types regulate both abundance of TAX1BP1 and its association with pA20 as well as other known binding partners such as TRAF6 and TNIP1 to regulate NF-kB during immune responses (Medhavy *et al*, 2024; Matsushita *et al*, 2016) and whether TAX1BP1 meets A20 during signaling complexes or aggrephagy (Sarraf *et al*, 2020; Samie *et al*, 2018).

Although a reduction in A20 abundance is a known cause of autoimmunity (Adrianto *et al*, 2011; Wang *et al*, 2013, 2016; Aeschlimann *et al*, 2018b; Kadowaki *et al*, 2018; Yu *et al*, 2020; Takagi *et al*, 2017; Zhou *et al*, 2016) we cannot be certain that the variants identified in *TNFAIP3* and *TAX1BP1* explains the clinical phenotype of A.II.1, nor was this the objective of the study. Rather, we have shown that the enormous diversity of human alleles provides a powerful resource for identifying mechanisms even in pathways that have been thoroughly investigated using genetic deletion mouse models. Just as elucidation of pathogenic alleles of single genes identified in patients with simple Mendelian diseases has clarified the function of many proteins, human genetic variants in epistasis can provide important new insights into protein-protein interactions.

## MATERIALS AND METHODS

### Patients

The patients described are part of a larger cohort of primary antibody-deficiency kindreds enrolled and recruited through the Australian and New Zealand antibody deficiency allele study and Centre for Personalised Immunology. This study has been approved by human research ethics committees at ACT Health and Australian National University (protocol ETH.1.15.015) each institution and was conducted in accordance with the Declaration of Helsinki.

### Whole exome sequencing

The Illumina paired-end genomic DNA sample preparation kit (PE-102-1001, Illumina) was used for preparing the libraries including end repair, A-tailing, and ligation of the Illumina adaptors. Each sample was prepared with an index using the Illumina multiplexing sample preparation oligonucleotide kit (PE-400-1001, Illumina) and then pooled in batches of 6 in equimolar amounts before exome enrichment. The Illumina TruSeq exome kit (FC-121-1008, Illumina) was used to capture the human exome for each sample pool. Each 6-plex exome-enriched library was sequenced in 2 lanes of an Illumina HiSequation 2000 with version 2 chemistry as 100-bp paired-end reads.

Sequence reads were mapped to the GRCh37 assembly of the reference human genome using the default parameters of the Burrows-Wheeler Aligner (bio-bwa.sourceforge.net). Untrimmed reads were aligned by allowing a maximum of 2 sequence mismatches; reads with multiple mappings to the reference genome were discarded along with polymerase chain reaction duplicates. Sequence variants were identified with SAMtools (samtools.sourceforge.net) and annotated using Annovar (www.openbioinformatics.org).

### Genotyping and Sanger sequencing

Genomic DNA was isolated from saliva using Oragene-DNA OG-500 kit (DNAgenoTek) according to the manufacturer’s instructions. Genomic DNA was then amplified with primers via a thermocycler. The presence of candidate mutations was then confirmed by Sanger sequencing.

### Flow cytometry

Peripheral blood mononucleated cells (PBMCs) were isolated by density gradient centrifugation on Ficoll and maintained in complete RPMI containing 10% fetal bovine serum, 2 mM l-glutamine (Sigma-Aldrich), and 100 U/mL penicillin and streptomycin (Sigma-Aldrich). For flow cytometry, PBMCs were washed with FACS buffer containing 2% Fetal bovine serum and 1% of NaN_3_ in phosphate-buffered saline and maintained at 4°C. Antibodies are listed in supplementary material.

### Proliferation assay

Naive CD4^+^ T cells were sorted by BD FACS Aria™ III Cell Sorter. Enriched naive CD4^+^ cells were then stained using Cell Trace Violet (CTV) Cell Proliferation Kit (Invitrogen) according to the manufacturer’s instructions. Cells were cultured with anti-CD3/CD28 expansion beads (Miltenyi Biotec) for 4 or 6 days. Proliferation of cells was then analyzed by flow cytometry.

### Plasmablast induction

PBMCs were seeded with complete RPMI and were cultured with various combinations of IL-21 (50ng/ml) and CD40L (1μg/ml), IL-21 (50ng/ml) and CpG (2μM) or IL-21 (50ng/ml) and CD40L transfectants L-cells for 4 or 5 days at 37°C with 5% CO_2_.

### CRISPR/Cas9 editing

pX330 U6 chimeric DD CBh hSpCas9 plasmid was obtained from Addgene (www.addgene.org). Guide RNA containing target cutting site (CCG) was constructed. In order to introduce desired mutations, DNA repair template was also synthesized (approximately 100bp). Cells were then transfected with these nucleotides by either Neon™ Transfection System (Invitrogen) according to the manufacturer’s instructions. Cells were then incubated at 37°C for 24 hours in humidified with CO_2_ incubator. The following day single cells were sorted by BD FACS Aria™ III Cell Sorter with reporter gene expression such as GFP and mCherry and the single cells were cultured for 3-4 weeks. Sanger sequencing and western blot were conducted to confirm knockout of the gene of interest.

### Immunoblotting

Cells were cultured at 37°C and treated with TNF (Recombinant Human TNFa (BioLegend®, 570104), Phorbol 12-myristate 13-acetate (PMA) (Stemcell™, 74042)/Ionomycin (Stemcell™, 73722) or left unstimulated. Cell lysates were prepared in RIPA buffer with a protease inhibitor cocktail (Roche) and a Halt phosphatase inhibitor (Pierce Net); the lysates were subjected to sodium dodecyl sulfate polyacrylamide gel electrophoresis (PAGE). DC protein assay (Bio-Rad) was performed according to manufacturer’s instructions and equal amounts of proteins were loaded on 10% of polyacrylamide gel. The gel was wet transferred into Immun-Blot® PVDF Membrane (Bio-rad). Primary antibodies were added and incubated at 4°C overnight. Horseradish peroxide–conjugated secondary antibodies were detected with Clarity Western ECL Blotting Substrate (Bio-rad). Band intensity was quantitated with ChemiDoc MP Imaging System (Bio-rad).

### Co-Immunoprecipitation

HEK293T cells were co-transfected with MYC tagged A20 and FLAG tagged TAX1BP1 expressing vectors. Total cell lysates were prepared with lysis buffer containing 20 mM Tris [pH 7.5], 137 mM NaCl, 2 mM EDTA, and 1% Triton X with a protease inhibitor cocktail (Roche) and a Halt phosphatase inhibitor (Pierce Net). Lysate was spun down at 13000rpm for 15 minutes at 4°C. Supernatant was then rotated with anti-FLAG magnetic beads (Sigma M8823) at 4°C overnight and immunoprecipitated the proteins according to manufacturer’s instructions. The samples were eluted with 4X SDS loading buffer and boiled at 95°C for 3 minutes. The samples were then immunoblotted with anti MYC or anti FLAG antibodies.

### Deubiquitination assay

The OTU domains of human *TNFAIP3* (encoding 2-370 residues) were cloned between BamHI and XhoI sites in pGEX-6P-1 vector (GE Healthcare). GST-fused TNFAIP3 OTU protein was expressed in *E. coli* BL21 (BIOLINE) cells at 30°C for 12-16 hrs with 200 µM IPTG. Cells were lysed by sonication in 50 mM Tris (pH8.0), 100 mM NaCl, and 1 mM EDTA, followed by the addition of Triton-X100 (1% w/v). The lysates were cleared by centrifugation and then incubated with glutathione agarose (SIGMA). Precipitated complex was washed three times with lysis buffer, followed by washing with Prescision buffer (50 mM Tris [pH 7.5], 150 mM NaCl, 1 mM EDTA, and 1 mM DTT). A20^OTU^ protein was excised with Precision protease (GE Health) and collected from supernatant. The purified A20^OTU^ domain was tested for hydrolysis of K63 linked tetraubiquitin chains in a time course experiment *in vitro*. Purified A20^OTU^ proteins were incubated in 100 µl of DUB buffer (25 mM HEPES [pH 8.0], 5 mM DTT, 5 mM MgCl2) containing 2.5 µg of K48- or K63- linked polyubiquitin chain (#UC-230, #UC-330 from BostonBiochem). At the indicated time, 20 µl of reaction mixture was collected and enzymatic reaction was stopped by the addition of SDS sample buffer. Negative control was known mutation A20^H256A^, which contains an amino acid substitution of His256 with alanine.

### Luciferase reporter assay

The pNfκB-Luc plasmid (InvivoGen) was co-transfected with the TNFAIP3 plasmid and Renilla reporter gene vector into HEK293 cells and incubated. The Dual-Glo® Luciferase Assay System (Promega) was conducted according to the manufacturer’s instructions to measure Luciferase and Renilla activity by Spark® multimode microplate reader (TECAN).

### RNAseq and Microarray

Naïve B cells (CD19+, CD27-, CD10- and CD21^high^) were isolated by BD FACS Aria™ II Cell Sorter. After sorting, the cells were incubated with anti Ig F(ab)_2_(5ug/ml) (Jackson ImmunoResearch Laboratories inc.) overnight at 37°C with 5% CO_2_. The activated naïve B cells were then used for RNA extraction. RNAs were extracted by either trizol method or RNA kit method. The RNA pellets were then resuspended with RNase free water and stored at -80°C for overnight. Duplicate biological samples were prepared for the assay. The microarray was done in Affymetrix platform with gene array.

Naïve T cells (CD4+CD3+ CCR7+ CD45RA+) were isolated by BD FACS Aria™ II Cell Sorter. Then cells were incubated with anti-CD2, CD3, and CD28 for 4 hours. Duplicate biological samples were prepared for the assay. RNAs were extracted by either trizol method or RNA kit method. Sequences then were analysed by R language.

The RNA solution was then delivered to Ramaciotti Centre for Genomics (Sydney) for microarray analysis and RNAseq. RNA libraries for RNAseq were prepared using Nextera XT prep according to manufacturer’s instructions and NextSeq 500 was run. For microarray, biotin labelled cDNA was hybridised to an Affymetrix Human PICO 2.0 Gene Array (100 format) for 16 hours at 45°C at 60rpm using an Affymetrix hybridization oven. The arrays were washed according to manufacturer’s instructions on an Affymetrix FS450 fluidics station. Arrays were scanned on an Affymetrix Scanner GC3000 7G.

### Gene expression

Gene constructs were obtained from VectorBuilder. The vectors containing *TNFAIP3*, *MALT1*, *BCL10*, *TRAF6* and /or *TAX1BP1* were transfected into HEK293 with lipofectamine (Invitrogen). The vectors containing *TNFAIP3* and /or *TAX1BP1* were transfected into Raji or knockout cell lines by electroporation Neon™. After transfection, cells were then incubated for 36 to 48 hours at 37°C with 5% CO_2_.

## Supplementary materials

Supplementary Fig. 1. Structural model of A20

Supplementary Fig. 2. Flow cytometric analysis of members of both *TNFAIP3* kindreds

Supplementary Fig. 3. Flow cytometric analysis of plasmablast induction in vitro.

Supplementary Fig. 4. Flow cytometric analysis of CD69 expression by CD4+ T cells.

Supplementary Fig. 5. Transcriptome analysis of B and T cells.

Supplementary Fig. 6. *TAX1BP1* variant

Supplementary Fig. 7. CRISPR/Cas9 gene-edited Raji cells

Supplementary Fig. 8. MALT1-mediated A20 cleavage

## Funding

Supported by NHMRC Program Grant 1113577 (MCC); NHMRC Centre for Research Excellence Grant 1079648 (MCC), NHMRC project grant 1049760, Royal Society Wolfson Fellowship RSWF\R2\222004 (MCC) and Pryor Bequest (CEL, MCC).

## Author contributions

Conceptualization: CEL, MCC

Data curation: CEL

Formal analysis: CEL, MD, KH, MCC

Investigation: CEL, MD, RC, KH, VA, BM

Visualization: CEL, MD, KH, MCC

Funding acquisition: CEL, MCC

Project administration: MCC

Supervision: MCC

Writing – original draft: ECL, MD, RC, KH, VA, BM, MCC

Writing – review & editing: CEL, MCC

## Competing interests

Authors declare that they have no competing interests.

## Data and materials availability

RNASeq GEO accession number GSE288284 and GSE288283

## REFERENCES

Adrianto I, Wen F, Templeton A, Wiley G, King JB, Lessard CJ, Bates JS, Hu Y, Kelly JA, Kaufman KM, et al (2011) Association of a functional variant downstream of TNFAIP3 with systemic lupus erythematosus. Nature Genetics 43: 253–258

Aeschlimann FA, Batu ED, Canna SW, Go E, Gül A, Hoffmann P, Leavis HL, Ozen S, Schwartz DM, Stone DL, et al (2018a) A20 haploinsufficiency (HA20): clinical phenotypes and disease course of patients with a newly recognised NF-kB-mediated autoinflammatory disease. Ann Rheum Dis 77: 728

Aeschlimann FA, Batu ED, Canna SW, Go E, Gül A, Hoffmann P, Leavis HL, Ozen S, Schwartz DM, Stone DL, et al (2018b) A20 haploinsufficiency (HA20): clinical phenotypes and disease course of patients with a newly recognised NF-kB-mediated autoinflammatory disease. Ann Rheum Dis 77: 728

Bates JS, Lessard CJ, Leon JM, Nguyen T, Battiest LJ, Rodgers J, Kaufman KM, James JA, Gilkeson GS, Kelly JA, et al (2009) Meta-analysis and imputation identifies a 109|[thinsp]|kb risk haplotype spanning TNFAIP3 associated with lupus nephritis and hematologic manifestations. Genes and Immunity 10: 470–477

Boone DL, Turer EE, Lee EG, Ahmad R-C, Wheeler MT, Tsui C, Hurley P, Chien M, Chai S, Hitotsumatsu O, et al (2004) The ubiquitin-modifying enzyme A20 is required for termination of Toll-like receptor responses. Nat Immunol 5: 1052–1060

Bosanac I, Wertz IE, Pan B, Yu C, Kusam S, Lam C, Phu L, Phung Q, Maurer B, Arnott D, et al (2010) Ubiquitin Binding to A20 ZnF4 Is Required for Modulation of NF-κB Signaling. Mol Cell 40: 548–557

Catrysse L, Vereecke L, Beyaert R & Loo G van (2014) A20 in inflammation and autoimmunity. Trends in Immunology 35: 22–31

Cheng J, Novati G, Pan J, Bycroft C, Žemgulytė A, Applebaum T, Pritzel A, Wong LH, Zielinski M, Sargeant T, et al (2023) Accurate proteome-wide missense variant effect prediction with AlphaMissense. Science 381: eadg7492

Compagno M, Lim WK, Grunn A, Nandula SV, Brahmachary M, Shen Q, Bertoni F, Ponzoni M, Scandurra M, Califano A, et al (2009) Mutations of multiple genes cause deregulation of NF-κB in diffuse large B-cell lymphoma. Nature 459: 717–721

Coornaert B, Baens M, Haegman M, Sun L & Marynen P (2008) T cell antigen receptor stimulation induces MALT1 paracaspase–mediated cleavage of the NF-κB inhibitor A20. Nature Immunology 9: 263–271

De A, Dainichi T, Rathinam CV & Ghosh S (2014) The deubiquitinase activity of A20 is dispensable for NF-κB signaling. EMBO reports 15: 775–783

Graham RR, Cotsapas C, Davies L, Hackett R, Lessard CJ, Leon JM, Burtt NP, Guiducci C, Parkin M, Gates C, et al (2008) Genetic variants near TNFAIP3 on 6q23 are associated with systemic lupus erythematosus. Nat Genet 40: 1059–1061

Heyninck K, Valck DD, Berghe WV, Criekinge WV, Contreras R, Fiers W, Haegeman G & Beyaert R (1999) The zinc finger protein A20 inhibits TNF-induced NF-kappaB-dependent gene expression by interfering with an RIP- or TRAF2-mediated transactivation signal and directly binds to a novel NF-kappaB-inhibiting protein ABIN. The Journal of cell biology 145: 1471–1482

Hori T, Ohnishi H, Kadowaki T, Kawamoto N, Matsumoto H, Ohara O & Fukao T (2019) Autosomal dominant Hashimoto’s thyroiditis with a mutation in TNFAIP3. Clin Pediatr Endocrinol 28: 91–96

Hu S, Wang Y, Gong Y, Liu J, Li Y & Pan L (2018) Mechanistic Insights into Recognitions of Ubiquitin and Myosin VI by Autophagy Receptor TAX1BP1. J Mol Biol 430: 3283–3296

Hutti JE, Turk BE, Asara JM, Ma A, Cantley LC & Abbott DW (2007) IκB Kinase β Phosphorylates the K63 Deubiquitinase A20 To Cause Feedback Inhibition of the NF-κB Pathway. Mol Cell Biol 27: 7451–7461

Iha H, Peloponese J, Verstrepen L, Zapart G, Ikeda F, Smith CD, Starost MF, Yedavalli V, Heyninck K, Dikic I, et al (2008) Inflammatory cardiac valvulitis in TAX1BP1-deficient mice through selective NF-κB activation. EMBO J 27: 629–641

Kadowaki T, Ohnishi H, Kawamoto N, Hori T, Nishimura K, Kobayashi C, Shigemura T, Ogata S, Inoue Y, Kawai T, et al (2018) Haploinsufficiency of A20 causes autoinflammatory and autoimmune disorders. J Allergy Clin Immunol 141: 1485–1488.e11

Kanayama M, Inoue M, Danzaki K, Hammer G, He Y-W & Shinohara ML (2015) Autophagy enhances NFκB activity in specific tissue macrophages by sequestering A20 to boost antifungal immunity. Nat Commun 6: 5779

Komander D & Barford D (2007) Structure of the A20 OTU domain and mechanistic insights into deubiquitination. Biochem J 409: 77–85

Lee EG, Boone DL, Chai S, Libby SL, Chien M, Lodolce JP & Ma A (2000) Failure to regulate TNF-induced NF-kappaB and cell death responses in A20-deficient mice. Science 289: 2350–2354

Ling L & Goeddel DV (2000) T6BP, a TRAF6-interacting protein involved in IL-1 signaling. Proc Natl Acad Sci 97: 9567–9572

Lu TT, Onizawa M, Hammer GE, Turer EE, Yin Q, Damko E, Agelidis A, Shifrin N, Advincula R, Barrera J, et al (2013) Dimerization and Ubiquitin Mediated Recruitment of A20, a Complex Deubiquitinating Enzyme. Immunity 38: 896–905

Ma A & Malynn BA (2012) A20: linking a complex regulator of ubiquitylation to immunity and human disease. Nature Reviews Immunology 12: 774–785

Martens A, Priem D, Hoste E, Vetters J, Rennen S, Catrysse L, Voet S, Deelen L, Sze M, Vikkula H, et al (2020) Two distinct ubiquitin-binding motifs in A20 mediate its anti-inflammatory and cell-protective activities. Nat Immunol 21: 381–387

Matsushita N, Suzuki M, Ikebe E, Nagashima S, Inatome R, Asano K, Tanaka M, Matsushita M, Kondo E, Iha H, et al (2016) Regulation of B cell differentiation by the ubiquitin-binding protein TAX1BP1. Sci Rep 6: 31266

Medhavy A, Athanasopoulos V, Bassett K, He Y, Stanley M, Tuipulotu DE, Cappello J, Brown GJ, Gonzalez-Figueroa P, Turnbull C, et al (2024) A TNIP1-driven systemic autoimmune disorder with elevated IgG4. Nat Immunol 25: 1678–1691

Morriswood B, Ryzhakov G, Puri C, Arden SD, Roberts R, Dendrou C, Kendrick-Jones J & Buss F (2007) T6BP and NDP52 are myosin VI binding partners with potential roles in cytokine signalling and cell adhesion. J Cell Sci 120: 2574–2585

Musone SL, Taylor KE, Lu TT, Nititham J, Ferreira RC, Ortmann W, Shifrin N, Petri MA, Kamboh MI, Manzi S, et al (2008) Multiple polymorphisms in the TNFAIP3 region are independently associated with systemic lupus erythematosus. Nature Genetics 40: 1062–1064

O’Neill TJ, Seeholzer T, Gewies A, Gehring T, Giesert F, Hamp I, Graß C, Schmidt H, Kriegsmann K, Tofaute MJ, et al (2021) TRAF6 prevents fatal inflammation by homeostatic suppression of MALT1 protease. Sci Immunol 6: eabh2095

Onizawa M, Oshima S, Schulze-Topphoff U, Oses-Prieto JA, Lu T, Tavares R, Prodhomme T, Duong B, Whang MI, Advincula R, et al (2015) The ubiquitin-modifying enzyme A20 restricts ubiquitination of the kinase RIPK3 and protects cells from necroptosis. Nat Immunol 16: 618–627

Oughtred R, Rust J, Chang C, Breitkreutz B, Stark C, Willems A, Boucher L, Leung G, Kolas N, Zhang F, et al (2021) The BioGRID database: A comprehensive biomedical resource of curated protein, genetic, and chemical interactions. Protein Sci 30: 187–200

Ramos PS, Criswell LA, Moser KL, Comeau ME, Williams AH, Pajewski NM, Chung SA, Graham RR, Zidovetzki R, Kelly JA, et al (2011) A comprehensive analysis of shared loci between systemic lupus erythematosus (SLE) and sixteen autoimmune diseases reveals limited genetic overlap. PLoS Genetics 7: e1002406

Razani B, Whang MI, Kim FS, Nakamura MC, Sun X, Advincula R, Turnbaugh JA, Pendse M, Tanbun P, Achacoso P, et al (2020) Non-catalytic ubiquitin binding by A20 prevents psoriatic arthritis–like disease and inflammation. Nature Immunology 21: 422–433

Samie M, Lim J, Verschueren E, Baughman JM, Peng I, Wong A, Kwon Y, Senbabaoglu Y, Hackney JA, Keir M, et al (2018) Selective autophagy of the adaptor TRIF regulates innate inflammatory signaling. Nat Immunol 19: 246–254

Sarraf SA, Shah HV, Kanfer G, Pickrell AM, Holtzclaw LA, Ward ME & Youle RJ (2020) Loss of TAX1BP1-Directed Autophagy Results in Protein Aggregate Accumulation in the Brain. Mol Cell 80: 779–795.e10

Shembade N, Harhaj NS, Liebl DJ & Harhaj EW (2007) Essential role for TAX1BP1 in the termination of TNF-α-, IL-1- and LPS-mediated NF-κB and JNK signaling. EMBO J 26: 3910–3922

Shembade N, Harhaj NS, Parvatiyar K, Copeland NG, Jenkins NA, Matesic LE & Harhaj EW (2008) The E3 ligase Itch negatively regulates inflammatory signaling pathways by controlling the function of the ubiquitin-editing enzyme A20. Nature Immunology 9: 254–262

Shembade N, Ma A & Harhaj EW (2010) Inhibition of NF-κB Signaling by A20 Through Disruption of Ubiquitin Enzyme Complexes. Science 327: 1135–1139

Skaug B, Chen J, Du F, He J, Ma A & Chen ZJ (2011) Direct, Noncatalytic Mechanism of IKK Inhibition by A20. Mol Cell 44: 559–571

Szklarczyk D, Franceschini A, Wyder S, Forslund K, Heller D, Huerta-Cepas J, Simonovic M, Roth A, Santos A, Tsafou KP, et al (2015) STRING v10: protein–protein interaction networks, integrated over the tree of life. Nucleic Acids Res 43: D447–D452

Takagi M, Ogata S, Ueno H, Yoshida K, Yeh T, Hoshino A, Piao J, Yamashita M, Nanya M, Okano T, et al (2017) Haploinsufficiency of TNFAIP3 (A20) by germline mutation is involved in autoimmune lymphoproliferative syndrome. J Allergy Clin Immunol 139: 1914–1922

Tewari M, Wolf FW, Seldin MF, O’Shea KS, Dixit VM & Turka LA (1995) Lymphoid expression and regulation of A20, an inhibitor of programmed cell death. J Immunol 154: 1699–1706

Valck DD, Jin D-Y, Heyninck K, Craen MV de, Contreras R, Fiers W, Jeang K-T & Beyaert R (1999) The zinc finger protein A20 interacts with a novel anti-apoptotic protein which is cleaved by specific caspases. Oncogene 18: 4182–4190

Wang S, Wen F, Tessneer KL & Gaffney PM (2016) TALEN-mediated enhancer knockout influences TNFAIP3 gene expression and mimics a molecular phenotype associated with systemic lupus erythematosus. Genes Immun 17: 165–170

Wang S, Wen F, Wiley GB, Kinter MT & Gaffney PM (2013) An Enhancer Element Harboring Variants Associated with Systemic Lupus Erythematosus Engages the TNFAIP3 Promoter to Influence A20 Expression. PLoS Genet 9: e1003750

Wertz IE, Newton K, Seshasayee D, Kusam S, Lam C, Zhang J, Popovych N, Helgason E, Schoeffler A, Jeet S, et al (2015) Phosphorylation and linear ubiquitin direct A20 inhibition of inflammation. Nature 528: 370–375

Wertz IE, O’Rourke KM, Zhou H, Eby M, Aravind L, Seshagiri S, Wu P, Wiesmann C, Baker R, Boone DL, et al (2004) De-ubiquitination and ubiquitin ligase domains of A20 downregulate NF-kappaB signalling. Nature 430: 694–699

Whang MI, Tavares RM, Benjamin DI, Kattah MG, Advincula R, Nomura DK, Debnath J, Malynn BA & Ma A (2017) The Ubiquitin Binding Protein TAX1BP1 Mediates Autophagosome Induction and the Metabolic Transition of Activated T Cells. Immunity 46: 405–420

White J, Suklabaidya S, Vo MT, Choi YB & Harhaj EW (2023) Multifaceted roles of TAX1BP1 in autophagy. Autophagy 19: 44–53

Yu M-P, Xu X-S, Zhou Q, Deuitch N & Lu M-P (2020) Haploinsufficiency of A20 (HA20): updates on the genetics, phenotype, pathogenesis and treatment. World J Pediatr 16: 575–584

Zammit NW, Siggs OM, Gray PE, Horikawa K, Langley DB, Walters SN, Daley SR, Loetsch C, Warren J, Yap JY, et al (2019) Denisovan, modern human and mouse TNFAIP3 alleles tune A20 phosphorylation and immunity. Nature Immunology 20: 1299–1310

Zhang M, Wang Y, Gong X, Wang Y, Zhang Y, Tang Y, Zhou X, Liu H, Huang Y, Zhang J, et al (2024) Mechanistic insights into the interactions of TAX1BP1 with RB1CC1 and mammalian ATG8 family proteins. Proc Natl Acad Sci 121: e2315550121

Zhou Q, Wang H, Schwartz DM, Stoffels M, Park YH, Zhang Y, Yang D, Demirkaya E, Takeuchi M, Tsai WL, et al (2016) Loss-of-function mutations in TNFAIP3 leading to A20 haploinsufficiency cause an early-onset autoinflammatory disease. Nature Genetics 48: 67–73

